# Unified cross-modality integration and analysis of T-cell receptors and T-cell transcriptomes

**DOI:** 10.1101/2023.08.19.553790

**Authors:** Yicheng Gao, Kejing Dong, Yuli Gao, Xuan Jin, Qi Liu

## Abstract

Single-cell RNA sequencing and T-cell receptor sequencing (scRNA-seq and TCR-seq, respectively) technologies have emerged as powerful tools for investigating T-cell heterogeneity. However, the integrated analysis of gene expression profiles and TCR sequences remains a computational challenge. Herein, we present UniTCR, a unified framework designed for the cross-modality integration and analysis of TCRs and T-cell transcriptomes for a series of challenging tasks in computational immunology. By utilizing a dual-modality contrastive learning module and a single-modality preservation module to effectively embed each modality into a common latent space, UniTCR demonstrates versatility across various tasks, including single-modality analysis, modality gap analysis, epitope-TCR binding prediction and TCR profile cross-modality generation. Extensive evaluations conducted on multiple scRNA-seq/TCR-seq paired datasets showed the superior performance of UniTCR. Collectively, UniTCR is presented as a unified and extendable framework to tackle diverse T-cell-related downstream applications for exploring T-cell heterogeneity and enhancing the understanding of the diversity and complexity of the immune system.

## Main

The adaptive immune system is one of the most complex and vital parts of the human body’s defence system, and T cells, with their T-cell receptors (TCRs), are integral components of this intricate network^1, 2^. The highly diverse TCRs on the surface of each T-cell can recognize a wide array of antigens, enabling the body to respond to various pathogens and malignant cells^3, 4^. The recent advent of single-cell sequencing technologies has enabled high-throughput profiling of TCR sequences and the corresponding gene expression profiles of individual T cells, providing an unprecedented view into the inner workings of the adaptive immune system^5-7^. Benefiting from these technologies, the development of computational methods that fully harness their capabilities is urgently needed.

Despite the rapidly developed multiple-modality sequencing approaches for TCRs and T-cell transcriptomes, however, many existing methods in immunology research have primarily focused on single-modality analysis, specifically examining single-cell gene expression profiles and TCR sequence data separately. For gene expression analysis, widely used tools such as Seurat^8^ (in R) and scanpy^9^ (in Python) have been employed to perform annotations on single-cell RNA sequencing (scRNA-seq) data. For TCR sequence analysis, methods such as TCR-BERT^10^, TCRdist^11^, and GLIPH^12^ have been utilized to identify potential antigen-specific clusters. While single-modality approaches have been valuable for investigating biological functions, they actually analyse cells in a localized manner, potentially overlooking the critical information derived from other aspects of a cell as well as interesting relationships across modalities. Considering the complexity of biological cells and their interactions across different modalities, methods that globally integrate and analyse different cellular modalities can offer more insightful results.

In recent studies, several multimodality analysis methods, including CoNGA^13^, Tessa^14^, and mvTCR^15^, have been introduced. These methods have demonstrated the value of integrating single-cell profiles and TCR sequences, leading to novel discoveries and offering a more comprehensive view of T cells than conventional single-modality analysis techniques. Despite their notable contributions, these methods do not yet provide a systematic and extendable strategy for various T-cell-related downstream task analyses, including single-modality analysis, multimodality analysis, cross-modality generation, and other vital aspects. Therefore, the development of a unified framework that can optimally utilize multimodality data to uncover profound insights in immunology research is needed^16^.

Here, we introduce UniTCR, a unified framework designed for the cross-modality integration and analysis of TCRs and T-cell transcriptomes, thereby enabling the execution of various tasks within computational immunology. UniTCR encompasses a dual-modality contrastive learning module as well as a single-modality preservation module to obtain the gene expression profile embedding and TCR embedding for each cell. By utilizing such embeddings, UniTCR can tackle an array of tasks in computational immunology, including (1) single-modality analysis; (2) modality gap analysis; (3) epitope-TCR binding prediction; and (4) TCR profile cross-modality generation. Notably, the single-modality analysis process in UniTCR is different from the traditional single-modality scRNA-seq/TCR-seq analysis method, as the embeddings in UniTCR are designed to incorporate information from the other modality. The versatility of UniTCR allows it to adapt to these diverse tasks by simply adjusting the weights of different modules during the model training process. In extensive evaluations conducted on multiple scRNA-seq/TCR-seq paired datasets, UniTCR consistently demonstrated superior or competitive performance. Collectively, UniTCR is presented as a unified and extendable framework to tackle diverse T-cell-related downstream applications for exploring T-cell heterogeneity and enhancing the understanding of the diversity and complexity of the immune system.

## Results

### 1. Overview of UniTCR

UniTCR is presented as a novel and unified single-cell embedding method designed for the joint analysis of individual T-cell gene expression profiles and their corresponding TCR sequences. Utilizing contrastive learning techniques^17^, UniTCR effectively embeds both T-cell gene expression profiles and TCR sequences into a shared latent space. Notably, UniTCR introduces an innovative and unique perspective that sets it apart from conventional embedding-based learning methods, where a modality preservation module is carefully designed. This module aims to capture robust latent representations and prevent overfitting on small datasets, which is a departure from traditional contrastive learning approaches that primarily depend on the utility of large datasets (Fig. 1). The whole process of UniTCR begins by encoding the input TCR sequence using Atchley factors^18, 19^, which are then passed through a self-attention-based feature representation encoder to embed the TCRs into a low-dimensional space. Concurrently, gene expression profiles are also embedded into a low-dimensional space through the use of multilayer neural networks^20^ (**see the Methods section**). To preserve the modality information within the original spaces, we calculate cell-to-cell distance matrices based on the original gene expression profiles and TCR-to-TCR distance matrices using TCR-BERT^10^, a large-scale pretraining model designed to capture the semantic information in TCR sequences (**see the Methods**). Then, new embeddings of T-cell gene expression profiles and TCR sequences that incorporate information from each modality are obtained; this is achieved through an integration of our contrastive objective and modality preservation objective (Fig. 1). Subsequently, UniTCR opens up unified and extendable avenues for downstream applications for investigating T-cell mechanisms and immune responses. These applications include single-modality analysis, modality gap analysis, epitope-TCR binding prediction, and cross-modality generation. Each application offers a specific perspective for studying and understanding the intricacies of T cells (Fig. 1). (1) Single-Modality Analysis: UniTCR provides a unique and comprehensive view of T-cell gene expression profiles and TCR embeddings for downstream analyses, offering a detailed understanding beyond what conventional single-modality approaches provide. (2) Modality Gap Analysis: By analysing the modality gap^21^ between two modalities, i.e., the gene expression modality and the TCR modality, UniTCR is capable of identifying potentially functional T-cell clusters via outlier detection with a modality gap, thus exposing key relationships that might otherwise go unnoticed with a single modality. (3) Epitope-TCR Binding Prediction: UniTCR demonstrates superior performance in three distinct testing scenarios for epitope-TCR binding prediction, including majority testing, few-shot testing, and zero-shot testing, which were described in our former study^22^. This highlights the benefits of incorporating gene expression profile information when handling a variety of real-world challenges involving epitope-TCR binding prediction. (4) Cross-Modality Generation: UniTCR offers a novel direction for computational immunology studies by providing cross-modality generation, particularly for the generation of gene expression profiles from TCR sequences when single-cell gene expression sequencing is not available. By designing a prior network^23^ and a profile decoder, UniTCR demonstrates the feasibility of this cross-modality generation strategy, offering a transformative approach for exploring T-cell functionality and immune responses (**see the Methods section**).

**Fig. 1:**
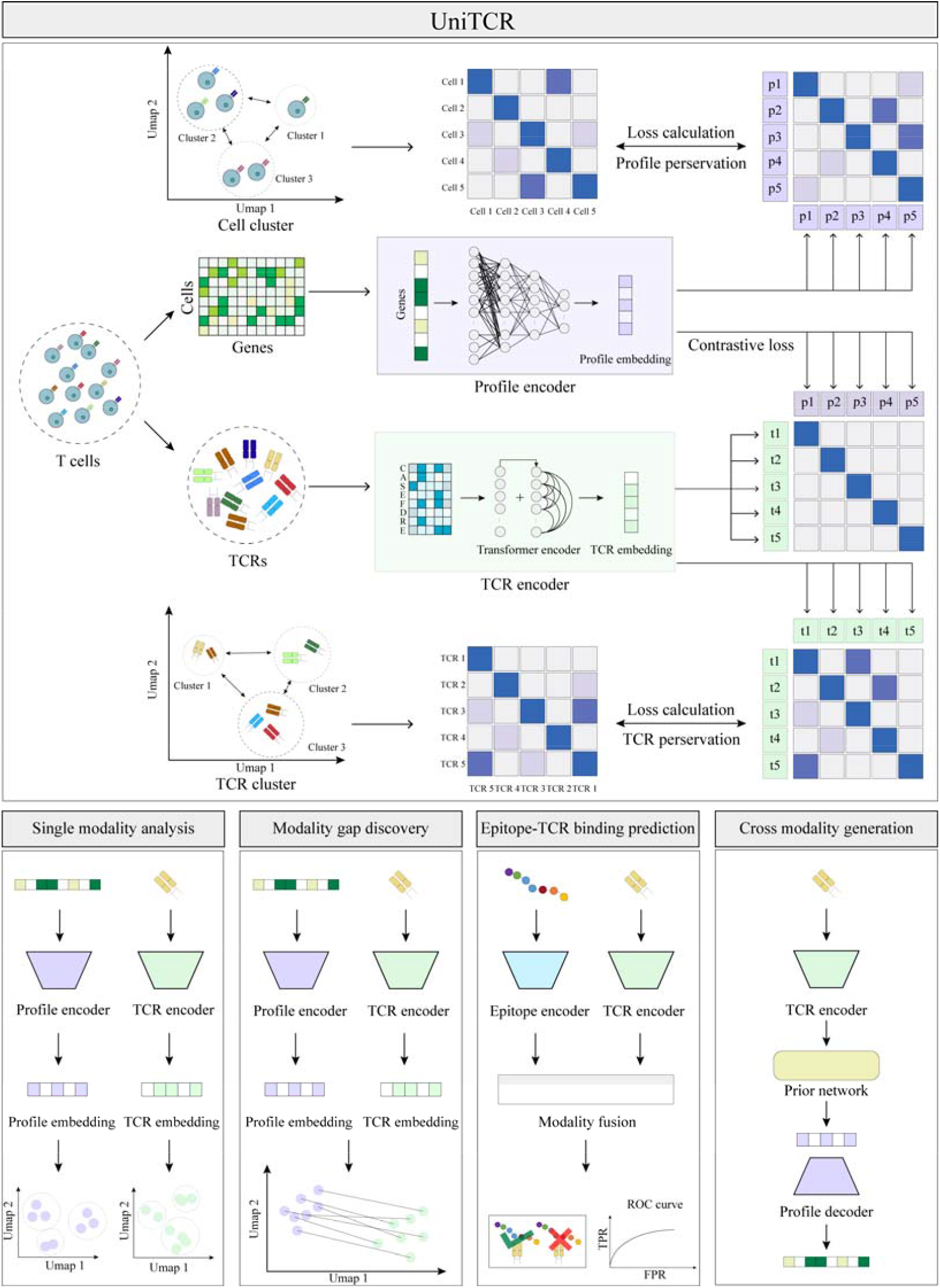
Illustration of the UniTCR framework. UniTCR consists of two modules.: a dual-modality contrastive learning module and a single-modality preservation module. UniTCR can be applied in four downstream applications. (1) Single Modality Analysis: The profile embedding/TCR embedding that incorporates information from the other modality is used for the downstream analysis. (2) Modality Gap Analysis: The modality gap between the profile embedding and TCR embedding is used to identify potentially functional cells. (3) Epitope-TCR Binding Prediction: The TCR encoder pretrained by the gene expression profile is used to construct an epitope-TCR binding prediction classifier. (4) Cross-Modality Generation: A prior network and a decoder network are constructed to generate gene expression profiles based on the pretrained TCR encoder.

### 2. Single-modality analysis - T-cell gene expression profile analysis with UniTCR

scRNA-seq is the predominant method for profiling T cells and identifying T-cell subpopulations and their corresponding functions^24^. UniTCR inputs normalized gene expression matrix and TCR sequences during the model training process. It maintains the intrinsic structures of different modalities by using the original cell-to-cell and TCR-to-TCR distance matrices. Subsequently, UniTCR generates low-dimensional embeddings of T-cell gene expression profiles by incorporating TCR information (**see the Methods section**).

As a result, we applied UniTCR to a peripheral blood mononuclear cell (PBMC) dataset from a patient who suffered kidney allograft rejection after anti-PD-1 therapy^25^ (denoted as the Kidney dataset) (Supplementary Note 2). To track and identify the pre-existing alloreactive T cells in the patient, a mixed-lymphocyte reaction (MLR) of recipient PBMCs and donor splenocytes was performed^25^. In a previous work, only the single-cell gene expression profile was used for cell clustering and annotation, and seven cell types were identified, including cytotoxic T lymphocyte (CTL) cells^26^, effector memory (EM) cells^27^, central memory (CM) cells^28^, mucosal-associated invariant T (MAIT) cells^29^, cycling cells^30^, Mito-hi cells^31^ and ZNF683+ cells^32^ (Fig. 2a). As a comparison, the profile embeddings derived from UniTCR, which incorporate TCR information, provided more detailed annotations of these cells while reliably maintaining the original cell type information (Figs. 2b, c and Supplementary Fig. 1a, b). Specifically, these clusters identified by UniTCR were categorized into clonotype-specific clusters and nonclonotype-specific clusters based on the percentage of identical clonotypes in each cluster (Fig. 2d; see the Methods section), where a dominant clonotype was observed in each clonotype-specific cluster. As an example, we focused on CTLs^26^, given that they represent the largest proportion of cells within these populations. Based on the proportion of CTLs within these clusters, the original CTL population in the reported study^25^ could be further divided into 13 clonotype-specific clusters and 2 nonclonotype-specific clusters by UniTCR (Fig. 2e, Supplementary Fig. 1c). By examining the activation^25^ and cytotoxic scores^33^ across these clusters, we found that different clonotype-specific clusters exhibited distinct biological functions, even though they were all CTLs (Supplementary Fig. 1e, f). Moreover, the T cells of varying clonotypes within the nonclonotype-specific clusters tended to exhibit similar biological functions, as expected (Supplementary Fig. 1g, h).

**Fig. 2:**
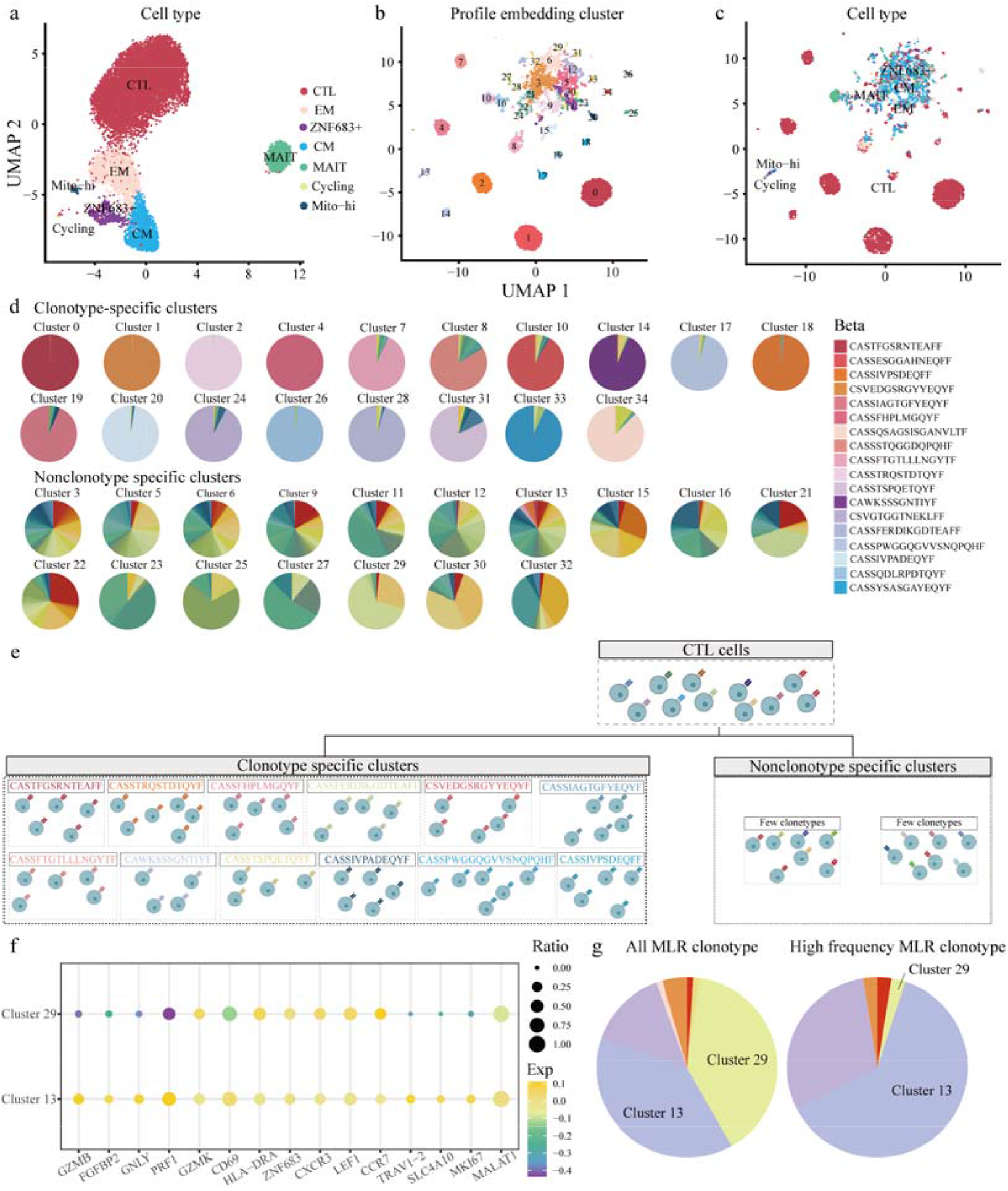
T-cell gene expression profile analysis results obtained with UniTCR. a. Uniform manifold approximation and projection (UMAP) clusters of T cells with their original gene expression profiles. b. UMAP clusters of T cells with the profile embeddings of UniTCR. c. UMAP clusters of T cells with the profile embeddings of UniTCR, annotated with the original cell types. d. The clusters identified by the UniTCR profile embeddings were categorized into clonotype-specific clusters and nonclonotype-specific clusters. e. The clusters identified by the UniTCR profile embeddings were further categorized into clonotype-specific clusters and nonclonotype-specific clusters for the original CTLs. f. The selected differentially expressed genes between cluster 29 and cluster 13. g. The ratio of MLR clonotypes and high-frequency MLR clonotypes in all clusters.

Our study also focused on ZNF683+ cell types, which were highlighted in previous research^25^ due to their potential role as alloreactive T cells. Characterized by high CXCR3, ZNF683 and HLA-DRA expression, these cells were primarily split into two clusters in the UniTCR profile embeddings (Supplementary Fig. 1d). Interestingly, these clusters showed different expression levels for other key marker genes and demonstrated diverse biological functions in terms of cell activation^25^ and cytotoxicity^33^ that go unnoticed with a single modality in the original study^25^ (Fig. 2f, Supplementary Fig. 1i, j). Furthermore, we analysed the potential alloreactive clonotypes, as identified by their MLR experiments. Although these clonotypes were primarily found in both clusters, the high-frequency clonotypes in the original study were particularly prevalent in cluster 13 (Fig. 2g).

Taken together, UniTCR demonstrates a remarkable capability to deliver a comprehensive perspective for T-cell analysis when provided with paired scRNA-seq/TCR-seq data. By incorporating TCR information, UniTCR is able to present fine-grained investigations that extend beyond what can be achieved using only single-cell gene expression data.

### 3. Single-modality analysis - T-cell receptor sequence analysis with UniTCR

Deep TCR sequence data analysis is vital when researchers are unsure about the specific antigen to which a TCR might bind. A common goal in such a study is to identify groups of TCRs sharing similar sequences, as these TCRs could potentially bind to the same antigen and assist in uncovering the potential physical and chemical properties of clusters. In the past, several TCR clustering methods were proposed to investigate epitope-specific T-cell responses^10-12^. However, these methods exclusively rely on the intrinsic information derived from TCR sequences. Notably, T-cell responses are also influenced by factors such as gene expressions and cellular states^34-36^. Therefore, a comprehensive analysis method that can incorporate these additional levels of gene expression information is expected to offer clearer insights.

To this end, UniTCR is designed to process input TCR sequences and generate TCR embeddings that incorporate the corresponding T-cell gene expression profile information. As a demonstration, we employed UniTCR on datasets derived from four 10x Genomics donors annotated with 44 epitope binding annotations. Then, the TCR embeddings were clustered, and the physicochemical properties, including the isoelectric point (PI), hydrophobicity, instability index (instaindex) and ratio between mass and charge number of ions (m/z), of each cluster were analysed^37^. Our results clearly showed that different clusters have different physicochemical properties (Fig. 3a, Supplementary Fig. 2a, 3a and 4a). Then, we used UMAP to visualize the TCR embedding for each donor, colour-coding it according to the three most prevalent epitopes in each donor (Figs. 3 b-g, Supplementary Fig. 2b-g, 3b-g and 4b-g). To assess the efficacy of UniTCR, we compared its TCR embeddings with those of TCR-BERT^10^, a pretraining model based on a large TCR sequence repertoire. Ideally, all clonotypes for each epitope should be closely embedded, which we quantified using the TCR compactness score (**see the Methods section**). As a result, our findings indicated that by incorporating profile information, UniTCR could yield superior TCR embeddings that are conducive to downstream TCR sequence analysis (Figs. 3 h-j, Supplementary Fig. 2h-j, 3h-j and 4h-j).

**Fig. 3:**
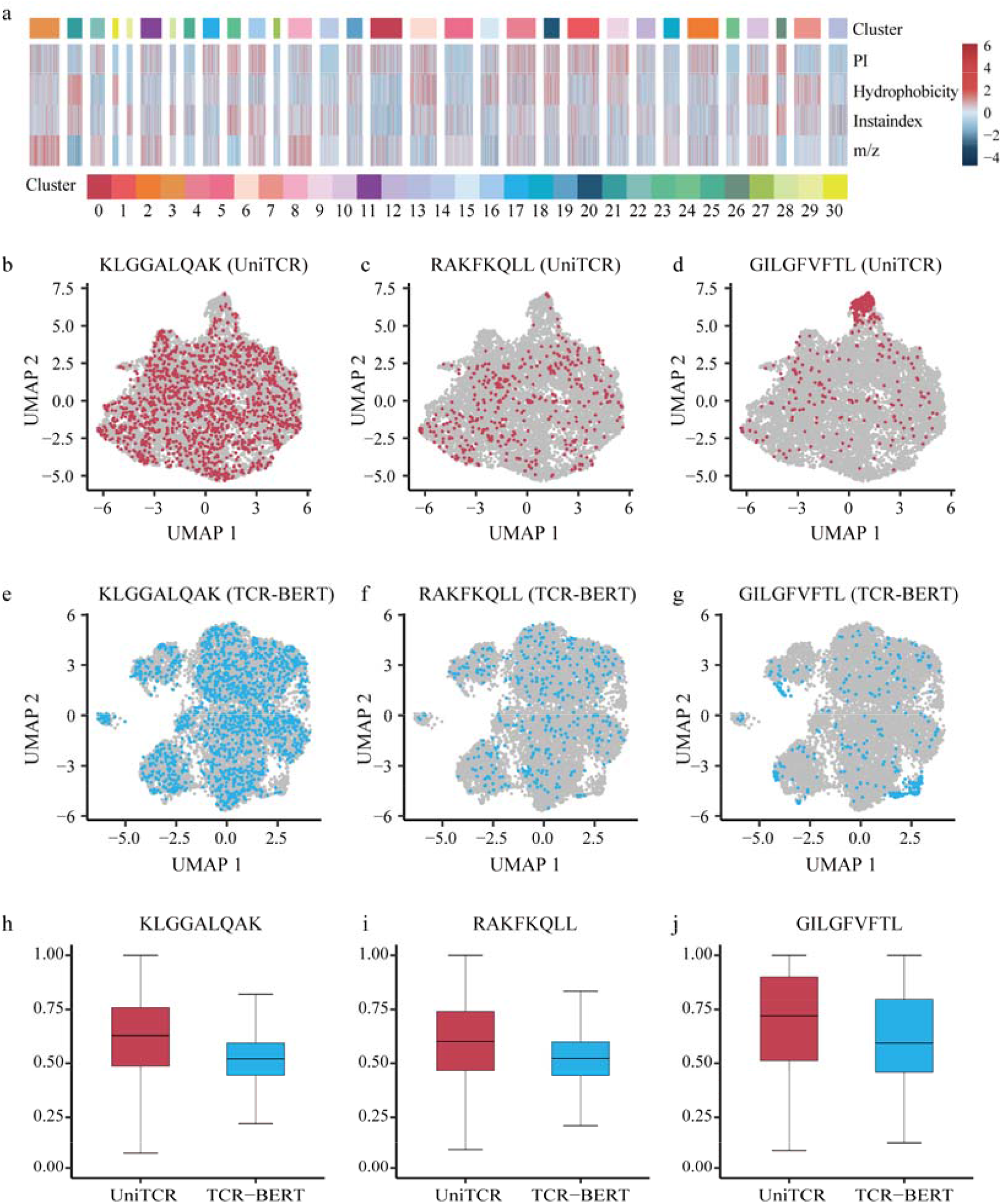
T-cell sequence analysis results obtained with UniTCR. a. Heatmap of the physicochemical properties of the TCRs in donor 2 clustered by the TCR embeddings of UniTCR. b. UMAP of the UniTCR TCR embeddings annotated by KLGGALQAK epitope specificity in donor 2. c. UMAP of the UniTCR TCR embeddings annotated by RAKFKQLL epitope specificity in donor 2. d. UMAP of the UniTCR TCR embeddings annotated by GILGFVFTL epitope specificity in donor 2. e. UMAP of the TCR-BERT TCR embeddings annotated by KLGGALQAK epitope specificity in donor 2. f. UMAP of the TCR-BERT TCR embeddings annotated by RAKFKQLL epitope specificity in donor 2. g. UMAP of the TCR-BERT TCR embeddings annotated by GILGFVFTL epitope specificity in donor 2. h. The TCR compactness scores between UniTCR and TCR-BERT for the KLGGALQAK epitope in donor 2. i. The TCR compactness scores between UniTCR and TCR-BERT for the RAKFKQLL epitope in donor 2. j. The TCR compactness scores between UniTCR and TCR-BERT for the GILGFVFTL epitope in donor 2. For each boxplot, the box boundaries represent the interquartile range, the whiskers extend to the most extreme data point (no more than 1.5 times the interquartile range), the black line in the middle of the box represents the median.

### 4. Modality gap analysis with UniTCR

The identification of outliers is a prevalent practice in biological and medical research, as these outliers may signify rare or unusual phenotypes or responses^38-40^. These outliers could potentially pave the way for the discovery of novel phenomena or therapeutic targets^41^. However, existing studies primarily concentrate on identifying outliers through a single modality. For instance, researchers might identify rare or significant cell clusters using single-cell profiles or peptide-specific clonotypes with TCR sequences. However, no methods have been devised to identify outliers based on the misalignment of two modalities. UniTCR can be regarded as a multimodal model based on contrastive learning^17^. This model embodies a fascinating geometric attribute in the representation space, which is commonly referred to as the modality gap^21^. In a dual-encoder multimodal model, the representations of the two modalities are distinctly separated when the model is initialized and continue to maintain a certain distance even after optimization^21^. Earlier research has elucidated the critical role of mismatched data in the formation of the modality gap under a low model temperature^21^. Nevertheless, the impact of multimodal data misalignment on the extent of the modality gap remains to be comprehensively examined. To this end, we designed an experiment to emulate varying degrees of misalignment by reshuffling known TCR-gene expression profile pairs from the kidney dataset (Fig. 4a). We strictly partitioned the datasets at each reshuffling level into training and validation sets, on which UniTCR was then trained. The modality gaps were visualized using UMAP in each reshuffling scenario (Fig. 4b). The results indicated that the modality gap was smallest when no reshuffling was performed, and it expanded proportionally with the extent of reshuffling (Figs. 4c and d).

**Fig. 4:**
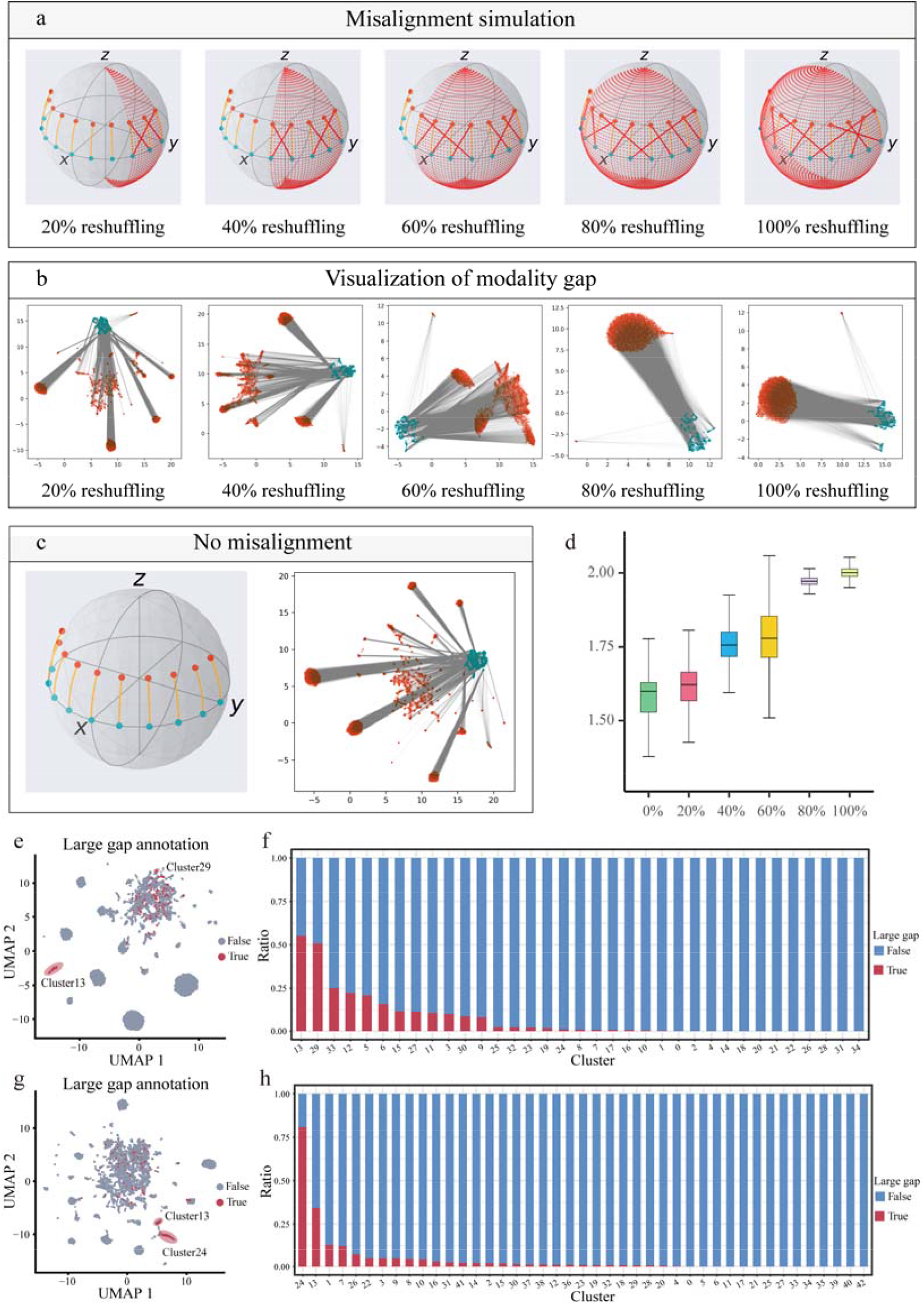
Modality gap analysis results obtained with UniTCR. a. Schematic diagram produced when simulating various degrees of misalignment by reshuffling the TCR-profile pairs in the Kidney dataset. b. UMAP of the profile embeddings and TCR embeddings in a common space, where blue represents TCRs, red represents gene expression profiles, and each black line connects the TCR embedding and profile embedding of the same cell. c. Schematic diagram of the case without misalignment in the Kidney dataset and the corresponding UMAP of the profile embeddings and TCR embeddings. d. The effect of the degree of misalignment on the modality gap. e. UMAP of the UniTCR profile embeddings annotated by the top 5% largest modality gaps in the Kidney dataset. f. The ratios of high-gap cells in different clusters for the Kidney dataset. g. UMAP of the UniTCR profile embeddings annotated by the top 5% largest modality gaps in the SCC dataset. h. The ratios of high-gap cells in different clusters for the SCC dataset. For each boxplot, the box boundaries represent the interquartile range, the whiskers extend to the most extreme data point (no more than 1.5 times the interquartile range), the black line in the middle of the box represents the median.

We then propose the hypothesis that the outliers among T cells, identified by the misalignment between the gene expression profiles and TCRs in the current cell population, may perform potentially crucial functions. Our reshuffling simulation suggests that a larger modality gap tends to correspond to a greater degree of misalignment between the gene expression profiles and TCRs of T cells. Therefore, we applied UniTCR to perform a modality gap analysis on the Kidney dataset^25^ (Fig. 4e). We classified the T cells with the top 5% largest modality gaps into the large-gap group and the rest into the small-gap group. Large-gap cells were predominantly found in clusters 13 and 29 (Fig. 4f). Interestingly, these clusters mainly consisted of the ZNF683+ cell type, which exhibits a high potential to have alloreactive T cells. We also discovered that the large-gap cells in this dataset generally exhibited increased activation and decreased cytotoxicity compared to the small-gap cells, which was consistent with the characteristics observed in clusters 13 and 29 (Supplementary Fig. 5a-d). We then applied UniTCR to the tumour-infiltrating lymphocyte (TIL) dataset derived from four squamous cell carcinoma (SCC) patients undergoing anti-PD-1 therapy^42^ (denoted as the SCC dataset) (Fig. 4g, Supplementary Note 3). Utilizing the UniTCR profile embeddings, we performed clustering on the T cells and calculated the modality gap for each cell. Interestingly, the large-gap cells were notably enriched with novel clonotypes and were found in clusters 13 and 24, with these clusters housing nearly 50% of such novel clonotypes and containing the clonotype with the highest frequency (Fig. 4h, Supplementary Fig. 6a-d). Additionally, we found that the large-gap cells demonstrated a higher exhaustion^33^ and proliferation score^33^ than the small-gap cells (Supplementary Fig. 6e, f). Notably, a previous study^42^ confirmed these novel clonotypes to be potentially tumour-specific, exhibiting a high exhaustion status (Supplementary Fig. 6 g, h). In our study, it was clearly shown that such novel clonotypes can be easily identified by modality gap analysis instead of experimental detections. Overall, there results indicated that investigating modality gap between T-cell receptors and T-cell transcriptomes is served as a useful and efficient indicator to identify T cells with potentially crucial functions.

**Fig. 5:**
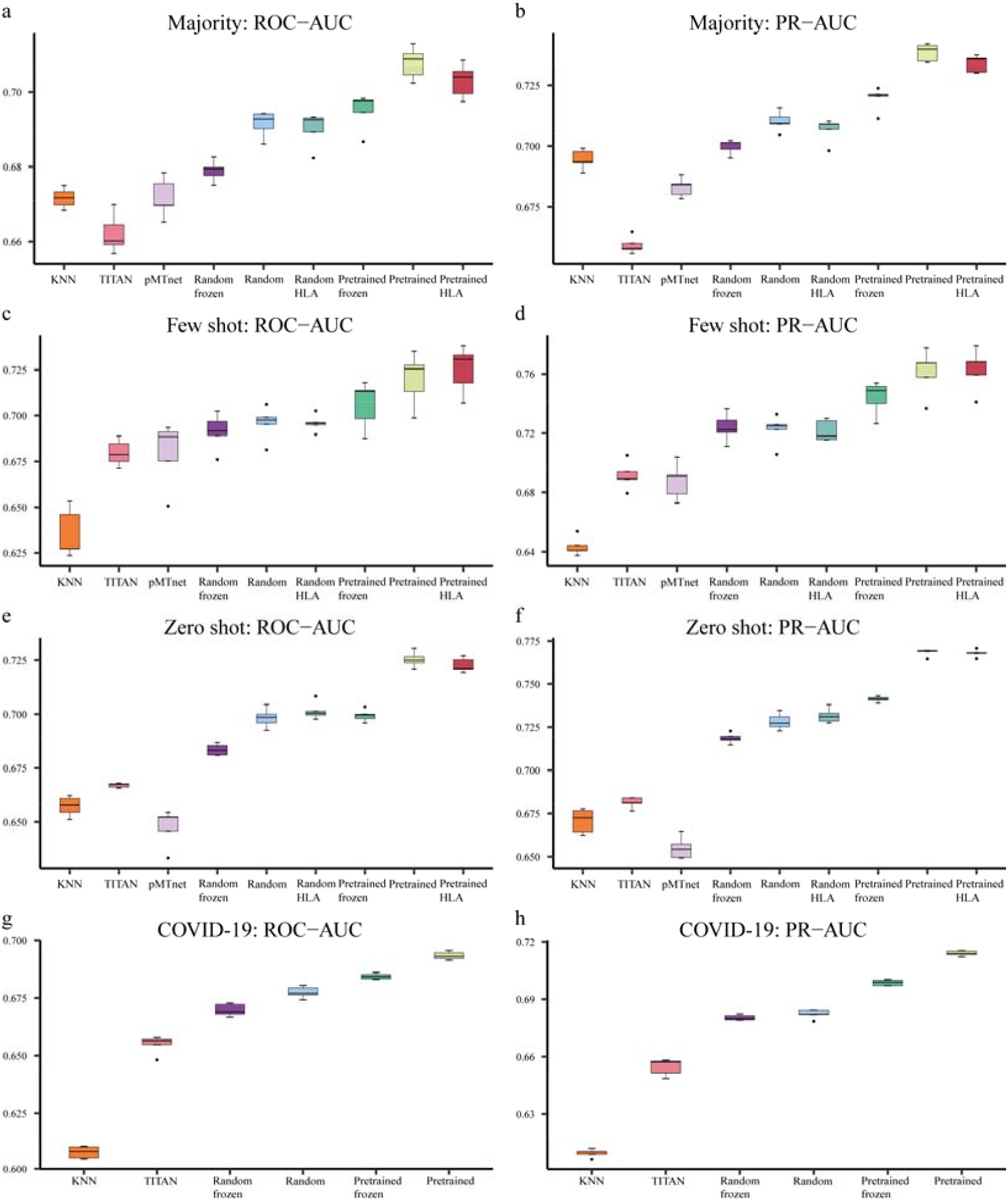
Epitope-TCR binding prediction task results obtained with UniTCR. a. The areas under the receiver operating characteristic curves (ROC-AUCs) of different classifiers in the majority testing setting. b. The areas under the precision-recall curves (PR-AUCs) of different classifiers in the majority testing setting. c. The ROC-AUCs of different classifiers in the few-shot testing setting. d. The PR-AUCs of different classifiers in the few-shot testing setting. e. The ROC-AUCs of different classifiers in the zero-shot testing setting. f. The PR-AUCs of different classifiers in the zero-shot testing setting. g. The ROC-AUCs of different classifiers for the independent COVID-19 dataset. h. The PR-AUCs of different classifiers for the independent COVID-19 dataset. All ROC-AUC and PR-AUC values were calculated with a random cross-validation split by five times. For each boxplot, the box boundaries represent the interquartile range, the whiskers extend to the most extreme data point (no more than 1.5 times the interquartile range), the black line in the middle of the box represents the median. KNN refers to the baseline model. Other abbreviations denote different initial settings under which UniTCR was trained. Pretrained: utilizing the pretrained TCR encoder for initialization. Pretrained frozen: freezing the pretrained TCR encoder. Random: initializing the TCR encoder with random values. Random frozen: freezing the TCR encoder with random initialization. Pretrained HLA: pretrained TCR encoder together with HLA. Random HLA: TCR encoder with random initialization and HLA.

**Fig. 6:**
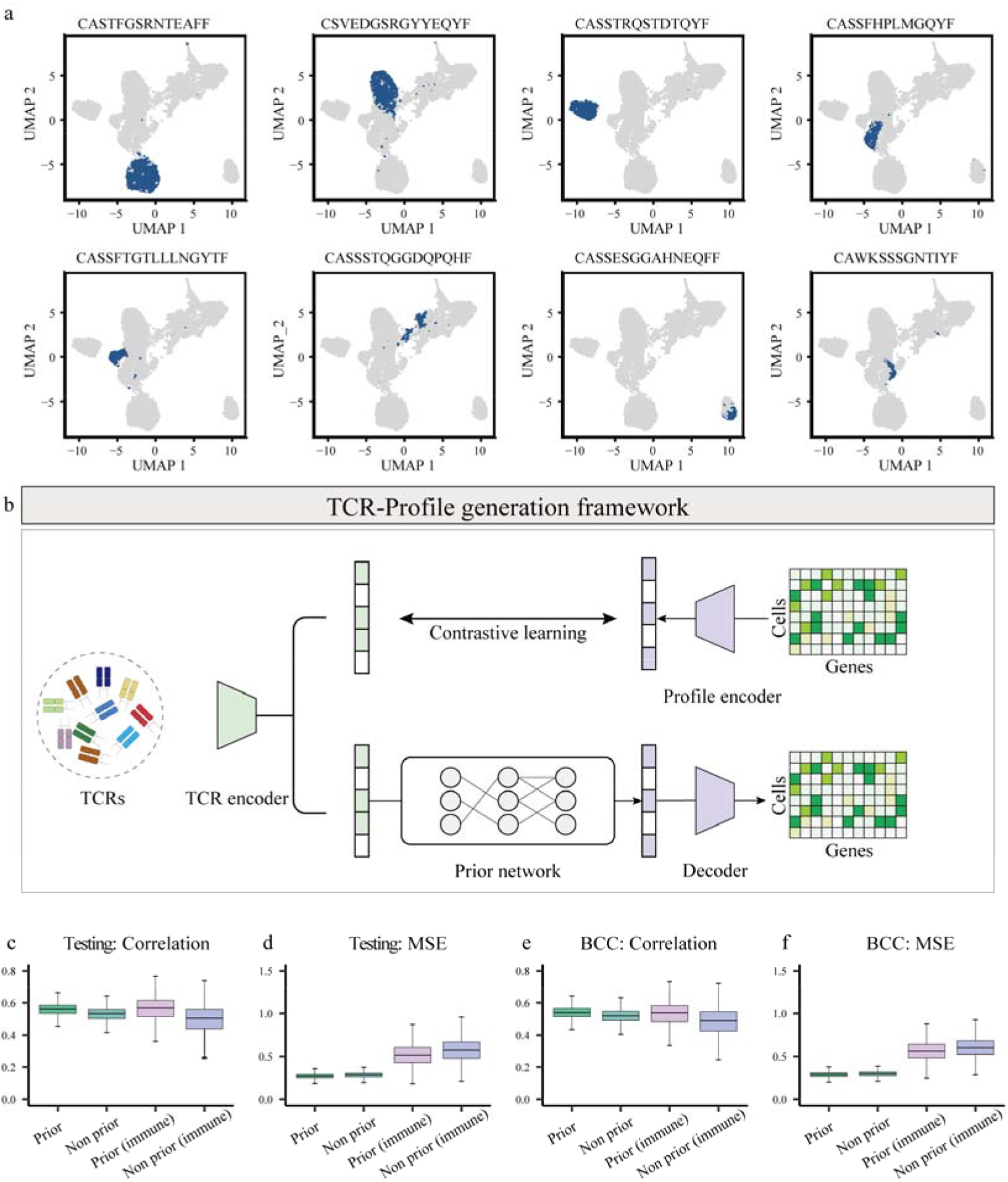
The cross-modality generation task results obtained with UniTCR. a. UMAP of T cells with the original gene expression profile annotated by the eight largest clonotypes. b. Illustration of the cross-modality generation framework. First, the TCR encoder was pretrained with a large scRNA-seq/TCR-seq paired dataset by contrastive learning. A prior network stacked with a gene expression profile decoder network was then constructed. Therefore, the TCR embeddings were generated by the TCR encoder, and the profile embeddings were predicted by the prior network. Finally, the gene expression profiles were decoded by the profile decoder. c. The Pearson correlation coefficients between the predicted and original gene expression profile produced with different settings for the testing dataset. d. The MSEs between the predicted and original gene expression profiles produced with different settings for the testing dataset. e. The Pearson correlation coefficients between the predicted and original gene expression profiles produced with different settings for the independent BCC dataset. f. The MSEs between the predicted and original gene expression profiles produced with different settings for the independent BCC dataset. For each boxplot, the box boundaries represent the interquartile range, the whiskers extend to the most extreme data point (no more than 1.5 times the interquartile range), the black line in the middle of the box represents the median. These abbreviations denote different settings. Prior: the model with the prior network, evaluated on all highly variable 5000 genes. No prior: the model without the prior network, evaluated on all highly variable 5000 genes. Prior (immune): the model with the prior network, evaluated on immune-related genes. No prior (immune): the model without the prior network, evaluated on immune-related genes.

Taken together, we clearly showed that the modality gap presented by UniTCR can not only be taken as the intriguing geometric phenomenon of the presentation of multimodal contrastive learning but also serve as a potentially useful biological discovery indicator for immunology studies.

### 5. Epitope-TCR binding prediction with UniTCR

The prediction of epitope-TCR binding specificity represents a formidable challenge in computational immunology, a task often equated to the field’s ‘holy grail’^43, 44^. The crux of the problem lies in the limited availability of data, which adheres to a long-tailed distribution^22, 43^. A small fraction of known epitopes have many associated TCRs, while the majority of epitopes are linked to a small number of TCRs or even lack any known TCR associations^22, 43^. This uneven data distribution imposes a substantial barrier to the accurate prediction of epitope-TCR bindings. Several general peptide-TCR binding prediction models have been proposed by embedding both peptides and TCRs^22, 45, 46^. However, constructing these models directly based on the currently available data may introduce a bias towards learning the binding patterns of epitopes with many known TCRs^22^. To circumvent this, our previous work^22^ suggested a more robust evaluation strategy using three distinct testing scenarios: majority testing, few-shot testing, and zero-shot testing, where majority testing refers to evaluating the performance of epitopes that have many known binding TCRs, few-shot testing refers to evaluating the performance for epitopes with only a handful of known binding TCRs and zero-shot testing refers to evaluating unseen epitopes that are not available in the training datasets (see the Methods section). This strategy allows for more accurate and unbiased predictions to be obtained across a range of epitope-TCR binding pairs.

As a demonstration, we first merged the Kidney dataset^25^, SCC dataset^42^, and the dataset from four 10x donors, as well as the SARS-CoV2 dataset^47^, with any batch effects removed. UniTCR was pretrained to acquire a TCR encoder that was capable of capturing profile information. Upon completion of the pretraining process, the TCR encoder from UniTCR was utilized as a foundation for constructing classifiers that predicted whether a specific TCR sequence could bind to a given antigen. In this context, UniTCR was treated as a black-box generator of TCR embedding vectors. Subsequently, an epitope encoder was designed to capture epitope sequence information, while a modality fusion encoder was developed to integrate both types of information (see the Methods section). Datasets sourced from IEDB^48^, VDJdb^49^, McPAS-TCR^50^ and PIRD^51^ were collected, and a subsequent quality control process was implemented (see the Methods section; Supplementary Table 1). This merged dataset was partitioned into zero-shot and nonzero-shot subsets based on the number of TCRs associated with each epitope (see the Methods section). The nonzero-shot dataset was further divided at a 3/1/1 ratio to assign the TCRs to training/validation/testing sets for each epitope. In addition, a large COVID-19 dataset^52^ was collected to serve as an independent set for evaluation purposes. To evaluate the efficacy of integrating profile information into TCR embeddings, random cross-validation was performed by five times for models with various initial settings (see the Methods section). We also compared UniTCR with other mainstream methods in this area, including k-nearest neighbours (KNN), pMTnet^46^ and TITAN^45^. Notably, PanPep^22^ was not included in this benchmark, as it was trained and tested in a pan-peptide meta-learning manner, thus the training/validation/testing datasets were different from those of the other methods. Then each model was evaluated in three testing scenarios and on the independent COVID-19 dataset, except for the comparison involving human leukocyte antigen (HLA) information (see the Methods section) (Figs. 6a-f). As a result, in comparison with all other methodologies, UniTCR with pretraining outperformed the competition, offering superior results. This robust performance is primarily attributable to the pretraining process of UniTCR, as it incorporated the gene expression profile information into the TCR encoder.

Taken together, our findings underscore the notion that the TCR encoder of UniTCR can serve as an effective and robust foundation for building classifiers that are capable of predicting epitope-TCR bindings. The integration of profile information can further boost the performance of these classifiers.

### 6. TCR profile cross-modality generation with UniTCR

With its rich diversity, the TCR repertoire offers insights into potential responses to a broad range of antigens^2^. Furthermore, the gene expression profile associated with each unique TCR sequence can provide detailed perspectives on the functional state of a T cell. However, due to its cost and labour intensiveness, single-cell sequencing remains a challenging process compared to bulk TCR sequencing, often rendering comprehensive T-cell analyses difficult^16^. Although various tools exist for calling TCR sequences from RNA-seq data^53-55^, efficient methodologies specifically designed to generate gene expression profiles from TCR sequences are lacking. The ability to predict T-cell gene expressions based on a TCR sequence holds significant potential to advance our understanding of immune responses. Cross-modality generation, which is instrumental in diverse applications such as image captioning, visual question answering, and multimodal translation, presents an innovative solution to this challenge^56, 57^. Our extensive analysis of TCR-profile pairs across different datasets revealed a consistent pattern; i.e., the same TCR clonotype tends to have similar gene expression profiles (Fig. 6a, Supplementary Fig. 7-9). This observation aligns with previous studies^13^, fuelling our motivation to develop a model that generates T-cell gene expression profiles from their respective TCR sequences.

To this end, UniTCR achieves cross-modality generation from TCRs to gene expression profiles by integrating a gene expression profile decoder with a prior (neural) network (see the Methods section). Previous research^23^ has indicated that models incorporating prior networks can weaken the impact of modality gap, exhibiting superior performance in cross-modality generation tasks. Accordingly, in our design, we incorporated a prior network to generate potential profile embeddings from given TCR embeddings. This approach was designed based on our observation of the existence of modality gaps between profiles and TCR embeddings, as described above, and such modality gap should be reduced before cross-modality generation. In this study, TCR embeddings were encoded by the TCR encoder, which was obtained through the contrastive learning process of UniTCR (Fig. 6b). To assess the performance of the model in this particular task, we merged datasets sourced from the Kidney dataset^25^, the SCC dataset^42^, four 10x Genomics donors, and the SARS-CoV2 dataset^47^ (see the Methods section). After removing batch effects, these datasets collectively yielded a total of 168,904 TCR-profile pairs. This integrated dataset was then divided into training and testing datasets at a ratio of 4:1. Additionally, we assembled an independent dataset derived from eleven patients with squamous cell carcinoma^42^ (BCC), denoted as the BCC dataset, comprising 14,490 TCR-profile pairs. The profile generation performance was evaluated by calculating the Pearson correlation coefficient^58, 59^ and the mean squared error (MSE)^60^ between the predicted and corresponding ground-truth expression values. Moreover, we computed the Pearson correlation coefficient and MSE between the predicted and ground-truth expression values of immune-related genes (Supplementary Table 2). A performance comparison was conducted between UniTCR and a variant of UniTCR without the prior network. Our results indicated that UniTCR, with the inclusion of the prior network, demonstrated superior performance across all testing scenarios (Fig. 6c).

## Discussion

The rapid development of single-cell high-throughput sequencing for profiling TCR sequences and T-cell transcriptomes has outpaced the corresponding computational framework required to gain integrative insights from such paired scRNA-seq/TCR-seq data. This highlights the need for a method that fully leverages the potential of multimodality data to uncover deeper insights in immunology research. Therefore, UniTCR is designed as a unified framework for integrating T-cell gene expression profiles with TCR sequences to address a wide array of computational immunology tasks. The findings and results obtained from our extensive evaluations demonstrate the potential of UniTCR in various contexts for immunology studies.

One of the most prominent features of UniTCR is its ability to perform single-modality analyses that incorporate information from another modality, providing a more holistic view than those yielded by conventional single-modality scRNA-seq or TCR-seq analyses. In addition, through our dual contrastive learning approach, we discovered that the geometric attributes of the modality gap could serve as meaningful indicators in this context. This finding provides a novel way to identify functionally relevant cells, which could have far-reaching implications for both basic research and therapeutic applications. In epitope-TCR binding prediction, UniTCR outperformed the existing state-of-the-art methods across multiple testing scenarios, further underlining the effectiveness of our integrative approach. Finally, UniTCR demonstrated a remarkable ability to generate gene expression profiles based on TCR sequences, which could greatly facilitate investigations into TCR-gene expression interactions.

Despite the encouraging results obtained in this study, we acknowledge that further improvements can be made. For instance, we did not consider the alpha chains of TCRs in our study as many previous works did^45, 46^, although encoding the alpha chains of TCRs can be an extensive module for UniTCR. Future updates expected when more paired scRNA-seq/beta chain/alpha chain data become available.

To our knowledge, UniTCR is a pioneering TCR profile cross-modality generation approach, offering a new direction for enhancing our understanding of T-cell functionality and immunity. While our current results are promising, the performance of this application can still be further enhanced by incorporating additional data in the future. Notably, due to the high cost associated with single-cell sequencing, the available datasets often exhibit significant biases towards TCRs with high specificity or those found in specific disease contexts^43^. The gene expression profiles of T cells are susceptible to changes based on different biological contexts. Therefore, to accurately capture the profile information associated with TCR sequences, there is a need for larger and more comprehensive datasets that cover a wider range of biological conditions.

In conclusion, the versatility of UniTCR facilitates its adaptation across various downstream research domains within immunology. The integration of TCR sequences and T-cell transcriptomes through UniTCR can elucidate the hidden interdependencies between these two modalities, thereby identifying functionally related T-cell clusters that might otherwise remain undetected in isolated analyses. Additionally, UniTCR can be adapted and tested on other types of multimodal data beyond scRNA-seq and TCR-seq data. This opens up a promising avenue for the comprehensive exploration of multimodal data in diverse biological contexts.

## Methods

### 1. Data curation

#### 1.1 Paired scRNA-seq/TCR-seq data

##### Data collection

The GEO database and 10X Genomics datasets were curated to gather pairwise scRNA-seq/TCR-seq data (Supplementary Table 1). Specifically, we collected four datasets from the GEO database, namely, the Kidney dataset (GSE216763)^25^, SCC dataset (GSE123813)^42^, SARS-CoV2 dataset (GSE191089)^47^ and BCC dataset (GSE123813)^42^. Additionally, we obtained paired data from four healthy donors in the 10X Genomics datasets. To ensure the quality of the data, the collected paired scRNA-seq/TCR-seq data were processed with a series of filtering steps (Supplementary Note 1). Only CD8+ T cells were retained, while genes expressed in fewer than three cells and cells expressing fewer than 200 genes or possessing mitochondrial genes expressing over 10% of the total expressed genes were filtered out. Cells were selected based on whether the beta chain was detected while ensuring that the length of the beta chain was in the range of 8-25. After filtering, we obtained paired data from the Kidney dataset with 11,894 cells, the SCC dataset with 11,162 cells, the SARS-CoV2 dataset with 10,244 cells, and the BCC dataset with 14,490 cells. Additionally, we acquired data from four healthy donors in the 10X Genomics datasets, consisting of 33,635, 61,583, 23,478, and 16,908 cells.

##### Batch effect removal

We used the Seurat package^8^ (version 3.2.2) to perform batch effect removal when integrating the pairwise scRNA-seq/TCR-seq data collected from various sources. The filtered data were first normalized using the NormalizeData function, and highly variable genes were identified using the FindVariableFeatures function with nfeatures = 5000. The batch effect was corrected using Seurat’s CCA method through three functions: SelectIntegrationFeatures, FindIntegrationAnchors and IntegrateData, where the nfeatures parameter in SelectIntegrationFeatures was set to 5000.

#### 1.2 Paired epitope-TCR data

##### Data collection

To acquire pairwise epitope-TCR data, five databases (EDB^48^, VDJdb^49^, McPAS-TCR^50^, PIRD^51^, and ImmuneCODE^52^) were utilized. Four of the databases (EDB^48^, VDJdb^49^, McPAS-TCR^50^ and PIRD^51^) provided comprehensive data for model training and testing, while the COVID-19 dataset extracted from the ImmuneCODE database, lacking HLA typing records, served as an independent dataset. During data collection, we focused on collecting HLA I-related epitope-TCR pairs, adhering to the following criteria: a) we explicitly included alpha chain, beta chain, HLA typing, and epitope information; b) HLA typing was specified in 2-field resolution; c) the alpha chain and beta chain sequences had lengths of 8 or greater, and the lengths of the epitope sequences fell within the range of 5-25 amino acids. Records from four of the databases, excluding COVID-19, needed to satisfy all three requirements to be retained. Finally, merging and deduplicating the data collected from IEDB, McPAS-TCR, VDJdb, and PIRD resulted in a collection of 22,273 unique epitope-TCR records, including 20,176 TCRs and 1,278 epitopes, denoted as the binding dataset. After removing any intersections with the aforementioned databases, the COVID-19 source provided an additional 471,363 independent epitope-TCR binding records associated with 511 epitopes.

##### Negative sample generation

Due to the absence of negative samples in the epitope-TCR records contained in the databases, we followed a similar approach to that of PanPep for negative sample generation^22^. Negative samples were generated by randomly sampling TCRs from a pool of healthy human PBMC TCRs^61^ for each peptide. The TCRs in this PBMC pools were filtered based on their lengths and amino acid compositions, and any overlaps with the four databases were removed. Subsequently, a subset of 20,176 records was randomly extracted from the processed data to serve as background data for model training. This number was equal to the total number of TCRs in the paired data collected from the four databases. The remaining data were randomly selected to match an equal number of negative samples for the validation and test datasets. Notably, the advantages of such a sampling strategy were demonstrated in a previous study^22, 62^.

##### HLA pseudosequence generation

In the epitope-TCR binding prediction application, we incorporated HLA subtype information, and we used pseudosequences to represent each HLA subtype. The pseudosequences were generated by following a methodology similar to that employed in netMHCpan^63^. First, the IPD-IMGT/HLA database^64^ was utilized to obtain the amino acid sequences for each subtype of human HLA. Subsequently, the HLA sequences were refined at the field 2 level, retaining only one record with the longest sequence among the HLAs with the same subtype. Additionally, HLA records with lengths of less than 171 amino acids were filtered out since the generation of pseudosequences requires that the given HLA sequence consist of at least 171 amino acids.

Finally, the HLA sequence positions mentioned in netMHCpan were extracted to derive the pseudosequences for the different HLA subtypes.

### 2. Design of UniTCR

We designed UniTCR and employed it in four application scenarios, namely, single-modality analysis, modality gap analysis, epitope-TCR binding prediction, and cross-modality TCR profile generation, to handle common and specialized joint analysis tasks involving paired TCR and T-cell gene expression profile data. The UniTCR framework comprises two core components: the dual-modality contrastive learning module and the single-modality preservation module. In the dual-modality contrastive learning module, the gene expression profiles and TCR sequences were embedded within a shared latent space. A profile encoder was carefully designed to learn a profile embedding that incorporates TCR information, while a TCR encoder was concurrently designed to learn a TCR embedding that incorporates profile information. In the single-modality preservation module, both the profile encoder and TCR encoder were trained to maintain the intrinsic relationships within each modality. By effectively leveraging these two components, UniTCR is equipped to handle the complexities of TCR and gene expression profile analysis, showcasing its utility in advanced immunological research.

Specifically, we consider a dataset containing *N* multimodal samples, each representing an individual T cell. Each sample *i* is composed of a gene expression profile, denoted by 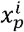, and a corresponding TCR sequence, denoted by 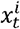, where *i* ∈ 1,…,*N* refers to the individual T cell. The gene expression profiles and TCR sequences are then grouped into batches of size *b*>1 based on their individual modalities, which results in 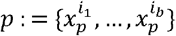 and 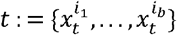. Our method treats these two input modalities separately, employing two carefully designed neural network encoders for each. The encoding for the gene expression profile is represented as 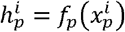, while the encoding for the TCR sequence is denoted as 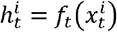. These encodings lead to the formation of *d*-dimensional vector representations,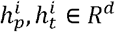, which are further processed by projection heads 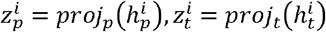, yielding 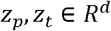. Each projection head is essentially a single linear multilayer perceptron (MLP) layer.

#### 2.1 Dual-modality contrastive learning module

In the dual-modality contrastive learning module, we utilize contrastive learning loss to align both the gene expression profile and TCR sequence modalities. The idea of this module is similar to that of the contrastive language-image pretraining (CLIP) method designed for cross-modality image and text analysis^17^. For the contrastive loss, we assume that we have *N* pairs of gene expression profiles and TCR sequences 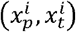, each with their respective representation 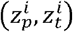. For the gene expression profile sample in the *i*^*th*^ pair, the corresponding TCR sequence 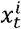 is considered the positive sample among the negative samples of the remaining individuals 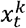 in the same batch. Similarly, for the TCR sequence sample 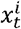, the gene expression profile 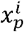 is regarded as the positive sample among the negative samples 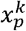 derived from the other individuals. Hence, the contrastive loss is the expectation of these two aspects: i) the profile-to-TCR *L*(*p,t*) and ii) TCR-to-profile *L*(*t,p*) aspects. Formally, during each training step, we randomly select a batch of size *b*>1 with indices {*i*_1_,…,*i*_*b*_} and employ the batchwise loss function as follows:

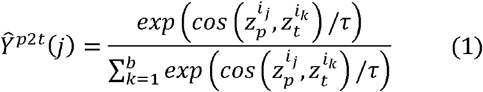

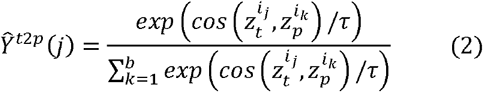

where *τ* is the model temperature parameter and *cos* is the cosine similarity. Let *Y*^*p*2*t*^(*j*),*Y*^*t*2*p*^(*j*) denote the ground-truth one-hot similarity, where negative pairs have probabilities of 0 and positive pairs have probabilities of 1. Then, the contrastive loss is defined as the cross-entropy *H* between *Y* and *Ŷ*:

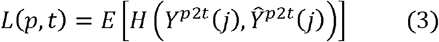

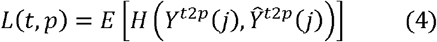

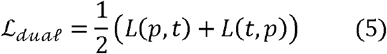

#### 2.2 Single-modality preservation module

Within the single-modality preservation module, we adopt distinctive loss objectives to conserve the inherent relationships that are present within each modality. For the gene expression profile modality, we compute cell-to-cell matrices using the Euclidean distance measure after performing normalization in the original gene expression profile space. For the TCR sequence modality, we leverage TCR-BERT to encode the TCR sequences, as this approach has been pretrained on a large TCR repertoire and can effectively capture the intrinsic semantic information embedded within the TCR sequences. The TCR sequences are then mapped into a 512-dimensional space. The TCR-to-TCR matrices are calculated based on cosine similarity after implementing L2 normalization for each TCR sequence. As a result, we design the preservation loss for each modality as follows:

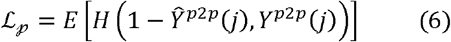

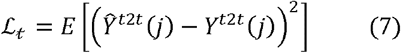

where *Y*^*p*2*p*^(*j*) is the element of the cell-to-cell matrix, and a lower value represents a closer relationship in the original space. *Y*^*t*2*t*^(*j*) ∈ [0,1] is an element of the TCR to TCR matrix, and a higher value signifies a closer relationship in the original space.

Therefore, the full pretraining objective of UniTCR is:

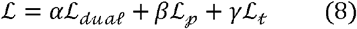

where *α+ β+ γ* = 1.

#### 2.3 Detailed model architecture of the single-modality encoder

UniTCR accepts gene expression profiles and TCR sequences as inputs, generating novel embeddings for each modality. Each modality employs a unique encoder to capture its inherent information. The gene expression profile encoder accepts an input containing 5,000 highly variable genes. The gene expression values are log-normalized and scaled. Subsequently, the encoder processes the input through three MLP layers with dimensions of 1,024, 512, and 256. The rectified linear unit (ReLU) function^65^ is employed as the activation function for these layers. Ultimately, layer normalization is applied to the encoder’s output. For the TCR sequence encoder, each TCR sequence is initially encoded into a 25x5 matrix via the Atchley factor^18^. In cases where the TCR sequence length falls short of 25, it is padded into the 25x5 configuration using a zero vector. The Atchley factor characterizes each amino acid with five numerical values, denoting its biochemical properties. Before learning from efficient semantic TCR information sequences, we employ the sinusoidal position encoding method. This approach embeds positional information into each amino acid within each TCR sequence. Additionally, we deploy a transformer encoder-based structure as the fundamental sequence encoder (Supplementary Fig. 10). The self-attention layer, an integral part of this structure, is a potent architecture comprising matrices *Q, K*, and *V*^66^. Accordingly, the input sequences, complemented with positional encoding, are initially processed by an embedding transformation layer and mapped into a 256-dimensional space. The embedding transformation layer includes a 256-dimensional MLP layer and a layer normalization layer, which is followed by a 0.15 dropout layer. The output obtained from the embedding transformation layer is then fed to the transformer encoder with four heads. The outputs from both encoders take the form of 256-dimensional embeddings. In the present study, UniTCR primarily focuses on the beta chains of TCRs due to their relative significance^22, 46^, while not taking the alpha chains of TCRs into account. However, it should be noted that our TCR encoder is highly adaptable and can also be readily extended to accommodate alpha chains.

### 3. Application of UniTCR

#### 3.1 Single-modality analysis with UniTCR

We applied UniTCR to conduct single-modality analyses on the Kidney dataset and the 10x datasets to illustrate that UniTCR can provide a more detailed view for analysing T cells. Utilizing normalized paired scRNA-seq/TCR-seq data as inputs, we generated profile embeddings from the Kidney dataset for profile embedding analysis, while the TCR embeddings from the 10x datasets were used for TCR embedding analysis purposes (Supplementary Note 2). For the profile embeddings, we normalized the data using the ScaleData of Seurat, focusing on 256 features. Principal component analysis (PCA) was performed on the normalized data using the RunPCA function, extracting 20 principal components. Cluster analysis followed, and it was performed using FindNeighbors and FindClusters with annoy.metric set to “cosine” and the resolution set to 0.5. For visualization, we applied UMAP projections to the selected principal components using RunUMAP. Clonotype-specific clusters were identified based on a threshold of 0.8, requiring 80% or more cells in a cluster to belong to the same clonotype. CTL-specific clusters were determined with a threshold of 0.7, as CTLs exhibited a distinct decreasing trend regarding the percentages of different clusters at this threshold. However, due to the low ZNF683+ cell count, the two clusters with the highest percentages of ZNF683+ cells were considered ZNF683+-specific. In the TCR embedding analysis, we deduplicated the data to ensure their uniqueness. Similar to profile embedding analysis, we conducted dimensionality reduction and clustering on the TCR embeddings. However, the parameter settings differed slightly, with the npcs parameter set to 50 for RunPCA and a resolution of 2 used for FindClusters. The Peptides package^37^ was employed to evaluate the physicochemical properties of the TCR sequences, including the PI, hydrophobicity, instability (InstaIndex), and mass-to-charge ratio (m/z). All analyses were conducted using R version 4.3.0 with default parameters unless otherwise specified. Additionally, the distribution of the TCRs targeting the same epitope in UMAP was examined. We compared the TCR embeddings of UniTCR with those of TCR-BERT, a large pretraining model based on the TCR sequence repertoire. The output of TCR-BERT was also processed with L2 normalization. We focused on the three epitopes with the largest numbers of known binding TCRs for each donor. The TCR compactness score served as an indicator to assess TCR dispersion, which is defined as follows:

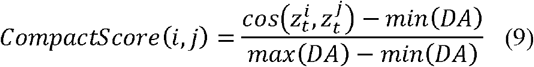

where TCR *i* and TCR *j* were associated with the same epitope, and *DA* is the cosine distance array of all TCR pairs for the donor.

#### 3.2 Modality gap analysis with UniTCR

Prior research^21^ has suggested that the contrastive learning objective may facilitate the occurrence of a modality gap. Consequently, in UniTCR, we define this modality gap as the difference between the TCR and profile embeddings for each individual cell. This can be expressed as:

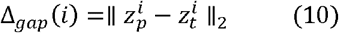

Here,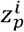 and 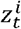 represent the L2-normalized gene expression profile and TCR sequence embedding, respectively, which are produced by their corresponding single-modality encoders in UniTCR. The Euclidean distance is an intuitive metric in this context, given that in UniTCR, both the profile and TCR embeddings are L2-normalized, signifying that they are always positioned on the unit sphere.

Explorations into the reasons behind the generation of modality gaps have indicated that the occurrence of “misaligned” data, particularly under low model temperatures, is a significant contributing factor, as suggested by previous studies. “Misalignment” implies that the ground-truth profile-TCR pairs should be 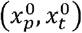 and 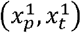, but the pairs obtained in reality are 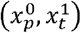 and 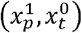. Nonetheless, previous studies have not examined the degree to which such misalignment in multimodal data might influence the modality gap. To address this, we designed a simulation experiment on the Kidney dataset. We introduced varying degrees of misalignment to this dataset by randomly selecting 20%, 40%, 60%, 80%, and 100% of the T cells with the paired profile and TCR sequences and performing reshuffling on the paired information for the cells we selected. This resulted in five simulated datasets. Each simulated dataset was then divided into training and validation datasets at a ratio of 4:1. Each training dataset was trained for 150 epochs using UniTCR, with the validation dataset used for early stopping. The weights of the different losses in UniTCR were 0.1, 0.8 and 0.1 in this setting. The AdamW optimizer^67^, with a learning rate of 0.0001, was used. Subsequently, we calculated the modality gaps for all T cells in each simulated training dataset.

#### 3.3 Epitope-TCR binding prediction

The task of epitope-TCR binding prediction can be constructed as learning a mapping function *Φ* from the space of TCR sequences *𝒯* and the space of epitopes *ε* to the space of binding outcomes.*ℬ*. This relationship can be formally defined as *Φ:𝒯*×*ε*→*ℬ*. The mapping function is learned using a training dataset 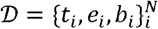, where *t*_*i*_ ∈ *𝒯,e*_*i*_ ∈ *ε* and *b*_*i*_ ∈ 0,1 is a binary label indicating whether a binding event has occurred.

Previous research has indicated that the evaluation of this task should be conducted in three more realistic scenarios: majority testing, few-shot testing, and zero-shot testing, due to the long-tailed distribution of the available data. ‘Majority testing’ refers to evaluating the performance of the model for epitopes that have many known binding TCRs, whereas ‘few-shot testing’ involves assessing the performance of the model for epitopes with only a handful of known binding TCRs. In the ‘zero-shot testing’ scenario, the epitopes are not included in the training datasets; instead, the model performance is evaluated on the testing dataset. Consequently, we partitioned our **binding dataset** by epitopes into two categories: a ‘nonzero dataset’ and a ‘zero dataset’. This division was based on whether an epitope had more than five available binding TCRs. The nonzero dataset was then subdivided into training, validation, and testing datasets, maintaining a ratio of 3:1:1 for each epitope. Subsequently, these training, validation, and testing sets ensured the consistency of the distributions across them. Epitopes with more than 100 known binding TCRs were used to evaluate the model’s performance in the majority testing scenarios. The remaining epitopes were utilized to evaluate its performance in few-shot testing scenarios. For the zero-shot testing scenario, we utilized both the zero-shot dataset and the COVID-19 dataset to assess our model’s performance. Notably, the epitopes in the COVID-19 dataset lack HLA information, rendering models that incorporate HLA sequence data unsuitable for evaluation on this dataset.

To evaluate UniTCR’s effectiveness in this crucial task, we devised an encoder specifically for epitopes. This encoder comprised a two-layer long short-term memory (LSTM) unit^68^ followed by a layer normalization layer (Supplementary Fig. 11). LSTM was chosen over a transformer-based architecture, similar to that used in the TCR encoder, because it exhibits superior inductive bias for sequence data, particularly when the available dataset is small. Each epitope 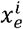 was first encoded by the Atchley factor and padded into a 25x5 matrix by the zero vector. Then, the epitope encoding, as the output of the epitope encoder, was represented as 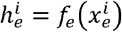. To better capture the interactions between the epitopes and TCRs, a cross-attention-based modality fusion block *f*_*F*_ was designed (Supplementary Fig. 11). The core multihead cross attention mechanism in UniTCR is defined as follows:

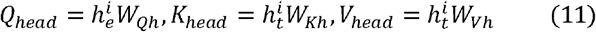

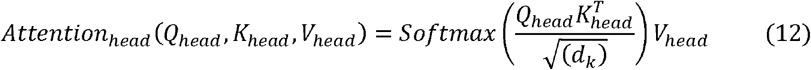

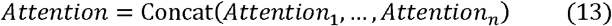

where 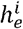 and 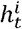 are epitope and TCR encodings from the epitope and TCR encoders, respectively. *W*_*Qh*_,*W*_*Kh*_ and *W*_*Vh*_ are the learnable weight matrices in one head. *d*_*k*_ is the scale factor and is equal to the number of heads in UniTCR, i.e., 4 heads. This mechanism allows the model to focus on different positions in parallel, improving the expressiveness of the attention mechanism. Hence, the binding prediction loss is defined as the cross-entropy *H* between *Y*_*et*_ and *Ŷ*_*et*_ :

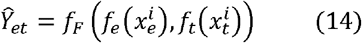

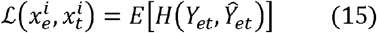

Here,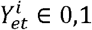 represents a binary label that signifies whether a binding interaction occurs between 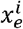 and 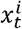 or not.

To assess the efficacy of incorporating gene expression profile information into the TCR encoder, we initially pretrained UniTCR on a large dataset, which included the Kidney, SCC, four 10x Genomics donors, and SARS-CoV2 datasets. To maximize the augmentation performance achieved for the gene expression profiles, we assigned the weights of the different loss components as 0.5, 0.5, and 0. The AdamW optimizer, with a learning rate of 0.0001, was used. UniTCR was pretrained for 200 epochs. The TCR encoder was then employed after post-pretraining as the foundation for constructing the classifier to predict the interaction bindings between the TCRs and the epitopes. To comprehensively assess UniTCR’s performance, we not only compared it with mainstream epitope-TCR binding prediction methods such as KNN, pMTnet, and TITAN but also explored various settings within UniTCR itself. These settings included utilizing the pretrained TCR encoder for initialization (Pretrained), freezing the pretrained TCR encoder (Pretrained frozen), initializing the TCR encoder with random values (Random), and freezing the TCR encoder with random initialization (Random_frozen). Additionally, models incorporating HLA information were constructed, such as a pretrained TCR encoder with HLA (Pretrained HLA) and a TCR encoder with random initialization and HLA (Random HLA). All these classifiers were trained and tested on the same datasets, and random cross-validation was performed five times. The zero-shot evaluation was performed on the zero-shot dataset and the COVID-19 dataset (except for the models incorporating HLA information). All classifiers underwent 250 epochs of training with the validation dataset used for early stopping. The AdamW optimizer, with a learning rate of 0.00001, was employed.

#### 3.4 Cross-modality generation

In this study, we focused on the cross-modality generation of TCRs to gene expression profiles because predicting (calling) a TCR sequence from RNA-Seq has been widely investigated. Therefore, the task of cross-modality generation in UniTCR can be constructed as learning a mapping function *Φ*_*c*_ from the space of TCR sequences *𝒯* to the space of gene expression profiles *𝒫*. This relationship can be formally defined as *Φ*_*c*_*:𝒯*→*𝒫*. The mapping function *Φ*_*c*_ is learned using a dataset 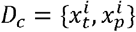, where 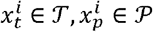. We then designed our generative stack to facilitate the generation of a gene expression profile from the given TCR sequence. This was accomplished using two main components: (1) a prior, denoted as 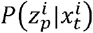, which generates a UniTCR gene expression profile embedding 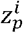 based on the given TCR,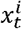; and (2) a decoder, represented as 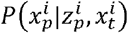, that yields a gene expression profile 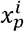 conditioned on the UniTCR gene expression profile embedding 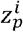 and the TCR 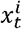. The decoder network enables the generation of a gene expression profile from its embedding, while the prior network facilitates the learning of a generative model for the profile embedding itself. Combining these two components leads to a generative model 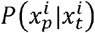 of the profile given the TCR sequence:

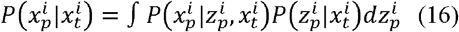

The prior (neural) network was designed for predicting profile embeddings based on TCR sequences. The TCR encoder that we pretrained in Methods Section 2.3 was used to produce the TCR embedding for each TCR. Then, we used a variational autoencoder^69^ (VAE)-based prior network to predict the profile embeddings from the TCR embeddings. Assume that the value of one profile embedding dimension comes from a Gaussian distribution. In this context, we learn the parameter of the Gaussian distribution *N*(*μ,σ*)from the TCR embedding generated from the TCR encoder, and the prior network produces new samples based on these learned parameters. The likelihood of observing a particular embedding value given a TCR embedding 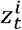 is defined as:

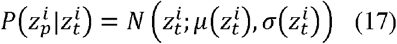

where 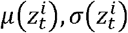are the middle outputs of the prior network, parameterized by *ϕ*_*p*_, given the TCR embedding 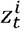. Then, the loss function of the prior network aims to maximize the following:

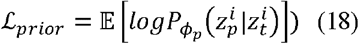

where 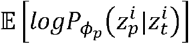 is the expected log-likelihood of the profile embedding data under the Gaussian model.

Although the zero-inflated negative binomial model has been widely used for capturing the distributions of count-based RNA-seq data^70, 71^, the RNA-seq data obtained after batch effect and removal lose their characteristics. Instead, our data exploration indicated that the gene value distribution is more likely to be the combination of two Gaussian distributions with,*N*(0,*σ*_1_) and N(*μ*_2_,*σ*_2_) which we defined as the *N*(0,*σ*)-inflated Gaussian distribution (Supplementary Fig. 12). Assuming that the value data of a gene come from an *N*(0,*σ*)-inflated Gaussian distribution, the associated probability density function (PDF) can be written as:

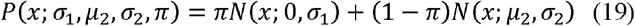

where *π* is the *N*(0,*σ*)-inflated parameter. *N*(0,*σ*) and *N(μ*_*2*_,*σ*_*2*_) denote the PDF of the Gaussian distribution with parameters 0,*σ*_1_ and,*μ*_2_,*σ*_2_ respectively. In the context of the decoder network, we learn the parameters of the -inflated Gaussian distribution (i.e.,*σ*_1_,*μ*_2_, *σ*_1_,*π*) from the data generated by the prior network. The decoder network can output these parameters as a function of the input data and produce new samples based on these learned parameters. To model the data using an *N*(0,*σ*)-inflated Gaussian distribution, the likelihood of observing a particular gene expression value given a latent variable 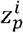 is defined as:

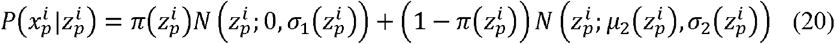

where 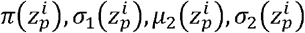 are the middle outputs of the decoder network, parameterized by *ϕ*_*d*_, given the gene expression profile embedding 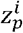. Then, the loss function of the decoder aims to maximize the following:

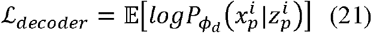

where 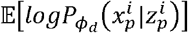 is the expected log-likelihood of the gene expression value data under the *N*(0,*σ*)-inflated Gaussian distribution model.

To evaluate the cross-modality generative model, we merged datasets from various sources, including Kidney, SCC, four donors from 10x Genomics, and SARS-CoV2. This merged dataset was subsequently divided into training and testing sets at a 4:1 ratio. Furthermore, we gathered a BCC dataset after removing batch effects to serve as an independent dataset for assessing the performance of the model. Our model evaluation not only employed the MSE and Pearson correlation metrics for all highly variable genes but also used these metrics specifically focused on immune-related genes (Supplementary Table 2). The AdamW optimizer, with a learning rate of 10e-6, was used. The model underwent a total of 200 epochs of training, with the first 70 epochs focused on the prior network. Following this, the prior network was frozen, and the next 130 epochs were dedicated to training the decoder network. The model without a prior network was also trained on the same model architectures but did not train the prior network via *ℒ*_*prior*_.

## Data availability

All datasets used by UniTCR for training, testing, and application examples have been made publicly accessible through the Github repository (https://github.com/bm2-lab/UniTCR). The paired scRNA-seq/TCR-seq data utilized for single modality analysis, modality gap analysis, and enhancing the prediction of TCR-pMHC binding were sourced from the GEO database (https://www.ncbi.nlm.nih.gov/geo/) and 10X Genomics Datasets. Specifically, the GSE accession numbers for the GEO datasets are as follows: GSE216763 (Kidnet dataset), GSE123813 (BCC dataset and SCC dataset), and GSE191089 (SARS-CoV2 dataset). Detailed information on the four healthy donors from the 10X Genomics Datasets is available at: https://www.10xgenomics.com/resources/datasets. For TCR-pMHC binding prediction model training and testing, data were obtained from databases including IEDB (https://www.iedb.org), VDJdb (https://vdjdb.cdr3.net), McPAS-TCR (http://friedmanlab.weizmann.ac.il/McPAS-TCR/), PIRD (https://db.cngb.org/pird/), and ImmuneCODE(https://clients.adaptivebiotech.com/pub/covid-2020). Additionally, control TCR data from 587 healthy donors were acquired through the original study (https://genomemedicine.biomedcentral.com/articles/10.1186/s13073-015-0238-z).

## Code availability

UniTCR is available on github (https://github.com/bm2-lab/UniTCR) together with a usage documentation and comprehensive example testing datasets.

## Acknowledgements

This work was supported by the National Key Research and Development Program of China (Grant No. 2021YFF1201200, No. 2021YFF1200900, received by Qi Liu), National Natural Science Foundation of China (Grant No. 31970638, 61572361, received by Qi Liu), Shanghai Shuguang scholars project (received by Qi Liu), Shanghai excellent academic leader project (received by Qi Liu), WeBank scholars project (received by Qi Liu) and Fundamental Research Funds for the Central Universities (received by Qi Liu).

## Author Contributions Statement

Qi Liu, Yicheng Gao and Kejing Dong designed the framework of this work. Yicheng Gao, Kejing Dong, Yuli Gao, Xuan Jin performed the analyses. Yicheng Gao, Kejing Dong and Qi Liu wrote the manuscript with the help of other authors. All authors read and approved the final manuscript.

